# <u>Heterotrophic bacterial diazotrophs</u> are more abundant than their cyanobacterial counterparts in metagenomes covering most of <u>the sunlit ocean</u>

**DOI:** 10.1101/2021.03.24.436778

**Authors:** Tom O. Delmont, Juan José Pierella Karlusich, Iva Veseli, Jessika Fuessel, A. Murat Eren, Rachel A. Foster, Chris Bowler, Patrick Wincker, Eric Pelletier

**Affiliations:** Génomique Métabolique, Genoscope, Institut François Jacob, CEA, CNRS, Univ Evry, Université Paris-Saclay, 91057 Evry, France; Research Federation for the study of Global Ocean systems ecology and evolution, FR2022/Tara GOsee, Paris, France; Institut de Biologie de l’ENS (IBENS), Département de biologie, École normale supérieure, CNRS, INSERM, Université PSL, 75005 Paris, France; Graduate Program in Biophysical Sciences, University of Chicago, Chicago, Illinois 60637, USA; Department of Medicine, University of Chicago, Chicago, Illinois 60637, USA; Bay Paul Center, Marine Biological Laboratory, Woods Hole, Massachusetts 02543, USA; Department of Ecology, Environment and Plant Sciences, Stockholm University Stockholm, 106 91, Sweden

**Keywords:** Marine bacteria, nitrogen fixation, open ocean, plankton, genomics, metagenomics, *Tara* Oceans, metagenome-assembled genomes

## Abstract

Biological nitrogen fixation is a major factor contributing to microbial primary productivity in the open ocean. The current view depicts a few cyanobacterial diazotrophs as the most relevant marine nitrogen fixers, whereas heterotrophic diazotrophs are more diverse and considered to have lower impacts on the nitrogen balance. Here, we used 891 *Tara* Oceans metagenomes to create a manually curated, non-redundant genomic database corresponding to free-living, as well as filamentous, colony-forming, particle-attached and symbiotic bacterial and archaeal populations occurring in the surface of five oceans and two seas. Notably, the database provided the genomic content of eight cyanobacterial diazotrophs including *Trichodesmium* populations and a newly discovered population similar to *Richelia*, as well as 40 heterotrophic bacterial diazotrophs organized into three main functional groups that considerably expand the known diversity of abundant marine nitrogen fixers compared to previous genomic surveys. Critically, these 48 populations may account for more than 90% of cells containing known *nifH* genes and occurring in the sunlit ocean, suggesting that the genomic characterization of the most abundant marine diazotrophs may be nearing completion. The newly identified heterotrophic bacterial diazotrophs are widespread, express their *nifH* genes *in situ*, and co-occur under nitrate-depleted conditions in large size fractions where they might form aggregates providing the low-oxygen microenvironments required for nitrogen fixation. Most significantly, we found heterotrophic bacterial diazotrophs to be more abundant than cyanobacterial diazotrophs in most metagenomes from the open oceans and seas. This large-scale environmental genomic survey emphasizes the considerable potential of heterotrophs in the marine nitrogen balance.

## Introduction

Plankton communities in the sunlit ocean consist of numerous microbial lineages that influence global biogeochemical cycles and climate^1–6^. Plankton primary productivity is often constrained by the amount of bioavailable nitrogen^7,8^, a critical element for cellular growth and division. Only a few bacterial and archaeal populations within the large pool of marine microbial lineages are capable of performing nitrogen fixation, providing a valuable source of new nitrogen to plankton^9–11^. These populations are known as diazotrophs and represent key marine players that sustain planktonic primary productivity in large oceanic regions^9^. Globally, marine nitrogen fixation is at least as important as the nitrogen fixation on land performed by *Rhizobium* bacteria in symbiosis with plants^12^.

Cyanobacterial diazotrophs are abundant in the surface of the open ocean and contribute to a substantial portion of nitrogen input^13–15^. They include populations within the genus *Trichodesmium*^16–18^ and other lineages that enter symbiotic associations with eukaryotes (e.g., *Richelia*^19,20^, the *Candidatus* Atelocyanobacterium also labeled UCYN-A^21,22^) or exist under the form of free living cells (some of the *Crocosphaera watsonii* cells also labeled UCYN-B^23,24^). A wide range of non-cyanobacterial diazotrophs has also been detected using amplicon surveys of the *nifH* gene required for nitrogen fixation. These molecular surveys showed non-cyanobacterial diazotrophs occurring in lower abundance compared to their cyanobacterial counterparts in various oceanic regions (e.g.,^25–32^) but could also be relatively abundant in some samples (e.g.,^33–37^). Overall, decades of *Trichodesmium* cultivation, flow-cytometry, molecular surveys, imaging and *in situ* nitrogen fixation rate measurements have led to the emergence of a view depicting cyanobacterial diazotrophs as the principal marine nitrogen fixers^38^.

Recently, a genome-resolved metagenomic survey exposed few free-living heterotopic bacterial diazotrophs (HBDs) abundant in the surface waters of large oceanic regions^39^. This first set of genome-resolved HBDs from the open ocean was subsequently found to express their *nifH* genes *in situ* using metatranscriptomics^40^. The metagenomic survey was focused on free-living bacterial cells, excluding not only key cyanobacterial players but also other diazotrophs that might occur under the form of aggregates, preventing a comprehensive investigation of diazotrophs in the sunlit ocean. To fill this gap, here we used nearly nine hundred *Tara* Oceans metagenomes^41^ to create a genomic database corresponding to free-living, as well as filamentous, colony-forming, particle-attached and symbiotic bacterial and archaeal populations occurring in surface waters of the global ocean. This database contains the genomic content of dozens of previously unknown HBDs abundant in different size fractions and oceanic regions, all found to express their *nifH* genes *in situ*. Most notably, we found HBDs to be more abundant (i.e., their genomic content was more represented) compared to cyanobacterial diazotrophs in metagenomes covering most of the surface of the open oceans and seas, emphasizing the considerable potential of heterotrophs in the marine nitrogen balance.

## Results and discussion

### Part one: Genome-wide metagenomic analyses

#### Nearly 2,000 manually curated bacterial and archaeal genomes from the 0.8-2,000 μm planktonic cellular size fractions in the surface oceans and seas

We performed a comprehensive genome-resolved metagenomic survey of bacterial and archaeal populations from polar, temperate, and tropical sunlit oceans using 798 metagenomes derived from the *Tara* Oceans expeditions. They correspond to the surface and deep chlorophyll maximum (DCM) layers from 143 stations covering the Pacific, Atlantic, Indian, Arctic, and Southern Oceans, as well as the Mediterranean and Red Seas, encompassing eight plankton size fractions ranging from 0.8 μm to 2000 μm (Table S1). These 280 billion reads were already used as inputs for 11 metagenomic co-assemblies using geographically bounded samples to recover eukaryotic metagenome-assembled genomes (MAGs)^42^. Here, we recovered nearly 2,000 bacterial and archaeal MAGs from these 11 co-assemblies.

Combining these MAGs with 673 MAGs previously generated from the 0.2 μm to 3 μm size fraction (93 metagenomes)^39^, we created a culture-independent, non-redundant (average nucleotide identity <98%) genomic database for microbial populations occurring in the sunlit ocean consisting of 1,778 bacterial and 110 archaeal MAGs, all exhibiting >70% completion (average completion of 87.1% and redundancy of 2.5%; Table S2). These 1,888 MAGs were manually characterized and curated using a holistic framework within anvi’o^43^ that relied heavily on differential coverage across metagenomes within the scope of their associated co-assembly. This genomic database has a total size of 4.8 Gbp, with MAGs affiliated to Proteobacteria (n=916), Bacteroidetes (n=314), Planctomycetes (n=154), Verrucomicrobia (n=128), Euryarchaeota (n=105), Actinobacteria (n=68), Cyanobacteria (n=51), Chloroflexi (n=36), Candidatus Marinimicrobia (n=30), Candidatus Dadabacteria (n=10) and 24 other phyla represented less than 10 times (Table S1). We used their distribution and gene content to survey marine diazotrophs in the open ocean without the need for cultivation or *nifH* amplicon surveys.

#### A genomic collection of 48 marine diazotrophs abundant in the open ocean

None of the 110 archaeal MAGs displayed signal for a diazotrophic life style. On the other hand, a total of 48 bacterial MAGs contained genes that encode the catalytic (*nifHDK*) and biosynthetic (*nifENB*) proteins required for nitrogen fixation (Table S3). Among them, the only absent gene was *nifH* missing in one MAG (Gammaproteobacteria), likely absent due to inherent limits of genome-resolved metagenomics. These diazotrophs could be categorized into eight cyanobacterial diazotrophs and 40 HBDs based on taxonomic signal confirmed by the occurrence of photosynthetic genes. Their estimated completion averaged 93.4%, suggesting they correspond to near-complete environmental genomes for dozens of diazotrophs abundant in different size fractions and oceanic regions (Figure 1 and Table S4).

**Figure 1:**
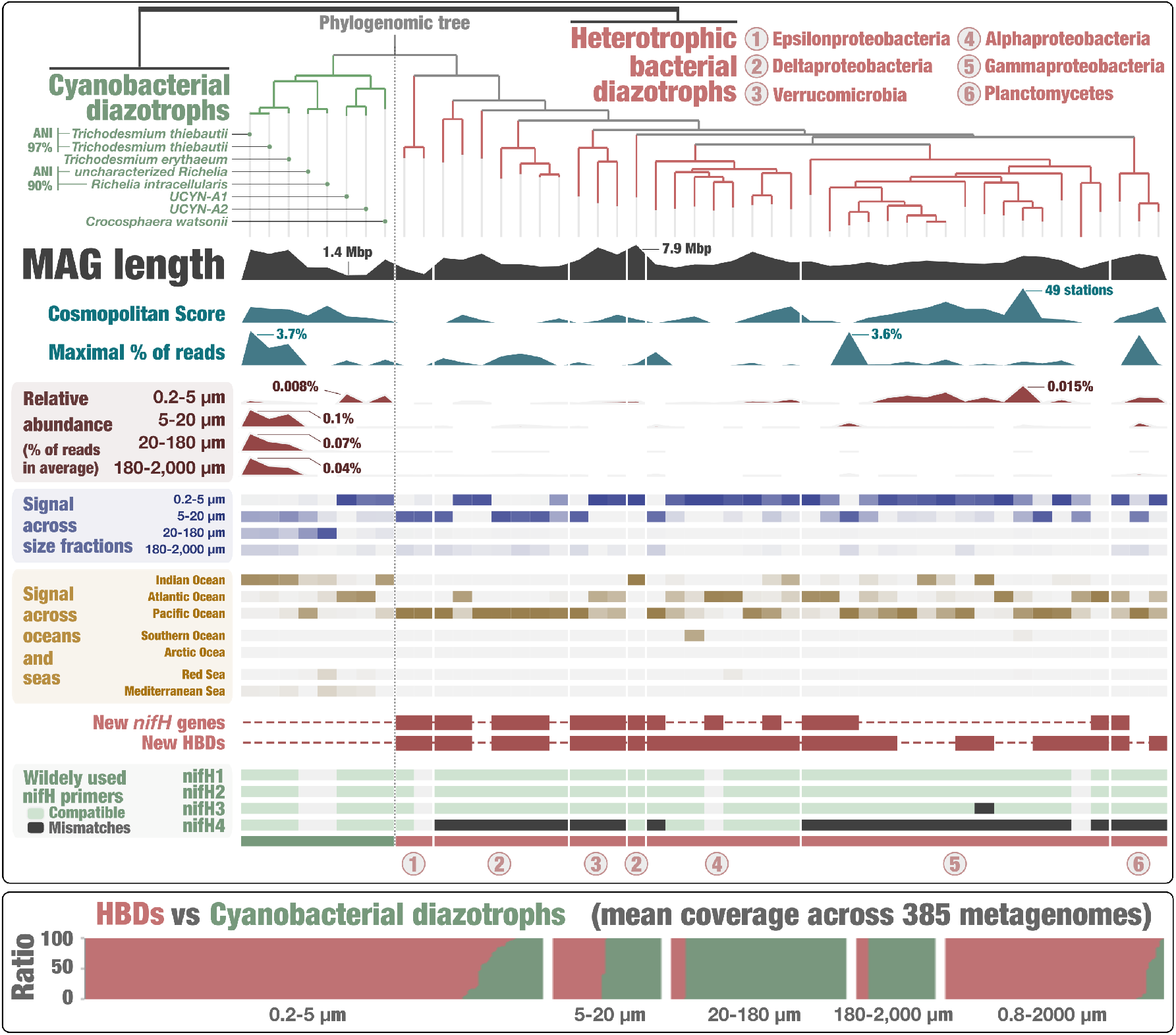
The phylogeny of 48 marine bacterial diazotrophs. Top panel displays a phylogenomic tree of the 48 diazotroph MAGs using 37 gene markers and visualized with anvi’o^43^. Additional layers of information display the length of MAGs alongside environmental signal computed using genome-wide metagenomic read recruitments across 937 metagenomes, and *nifH* primer compatibilities (only full length and non-fragmented *nifH* genes were considered). Bottom panel displays the ratio of cumulative genome-scale mean coverage between eight cyanobacterial diazotrophs (green) and 40 HBDs (red) across 385 metagenomes we organized into five size fractions.

Cyanobacterial MAGs recapitulated findings of major marine diazotrophs previously discovered within this phylum and for which a genome (partial or complete) has been characterized using either culture or flow cytometry: UCYN-A1 (ANI of 99.3%) and UCYN-A2 (ANI of 99.6%), *Crocosphaera watsonii* (strain WH-8501; ANI of 99.4%), *Richelia intracellularis* (strain RintHH01; ANI of 99.5%), *Trichodesmium erythraeum* (strain IMS101; ANI of 99%), and *Trichodesmium thiebautii* (strain H9-4; n=2 with ANI of 98.7% and 98%). Interestingly, while the two *Trichodesmium thiebautii* populations displayed high genomic similarity (ANI of 97.9%) and linear correlation across 81 metagenomes with signal (R^2^=0.93), mean coverage ratio revealed one dominant population in three sites of the North Atlantic Ocean while the other one occurred relatively more in the Indian Ocean, Pacific Ocean and Red Sea (Figure S1). In addition, one MAG with GC-content of 34.6% corresponded to a new population we tentatively named ‘*Candidatus* Richelia exalis’ given its close evolutionary relationship with *R. intracellularis* (e.g., ANI of 87.3% when compared to the strain RintHH01; see Table S3 for more comparisons) (Figure 1). The strong signal of ‘*Candidatus* Richelia exalis’ in the large size fractions, similar to *R. intracellularis*, and their comparable functions traits (see following section) suggests this species has also entered a symbiotic association within plankton.

Compared to the cyanobacterial diazotrophs already well characterized before this genome-resolved metagenomic survey, the HBDs we recovered substantially broaden the number of known diazotrophic populations abundant in the sunlit ocean. Aside from eight HBDs already characterized from the 0.2–3 μm size fraction^39^ (5 of which were replaced by MAGs characterized from the larger size fractions and displaying better completion statistics), the genomic database included 32 additional HBDs that not only expanded the known diversity of marine nitrogen fixers within Deltaproteobacteria (total of 8 HBDs with 6 new *nifH* genes when compared to a comprehensive set of reference databases^17^, see methods), Gammaproteobacteria (total of 16 HBDs with 4 new *nifH* genes) and Planctomycetes (total of 3 HBDs with 1 new *nifH* gene) but also covered populations within Alphaproteobacteria (n=8; three new *nifH* genes), Epsilonproteobacteria (n=2; all with new *nifH* genes), and Verrucomicrobia (n=3; all with new *nifH* genes) (Figure 1 and Table S5). Interestingly, some of the newly identified *nifH* gene sequences are incompatible with the design of some widely used primers (Figure S2 and Table S6). This was especially true of the “nifH4” primer (round one of the nested primers) many amplicon surveys are based upon^34,44–46^ (Figure 1), apparently incompatible with most HBDs abundant in the sunlit ocean.

#### The emergence of three main functional groups for marine HBDs

In order to provide a global view of functional capabilities among the 48 diazotrophs, we accessed functions in their gene content using COG20 functions, categories and pathways^47^, KOfam^48^, KEGG modules and classes^49^ from within the anvi’o genomic workflow^43^ (Table S7). Genomic clustering based on the completeness of 322 functional modules exposed four distinct groups: (1) the cyanobacterial diazotrophs, (2) HBDs dominated by Alphaproteobacteria, (3) HBDs solely from Gammaproteobacteria, and finally (4) HBDs organized in closely related subgroups corresponding to Deltaproteobacteria, Epsilonproteobacteria, Verrucomicrobia and Planctomycetes (Figure 2). Interestingly, one HBD (genus *Marinibacterium*) contained the *pufM* and *pufL* genes for anoxygenic photosystem II, denoting a photoheterotrophic life style. Furthermore, multiple HBDs contained functions relating to cobalamin biosynthesis (especially the Deltaproteobacteria) or denitrification (especially the Gammaproteobacteria), contributing to other important aspects of the planktonic productivity and nitrogen cycle (Table S7). Overall, we found a strong functional signal for the taxonomical lineages of marine diazotrophs discovered thus far, with the HBDs organized into three main groups and functionally more diverse compared to their cyanobacterial counterparts.

**Figure 2:**
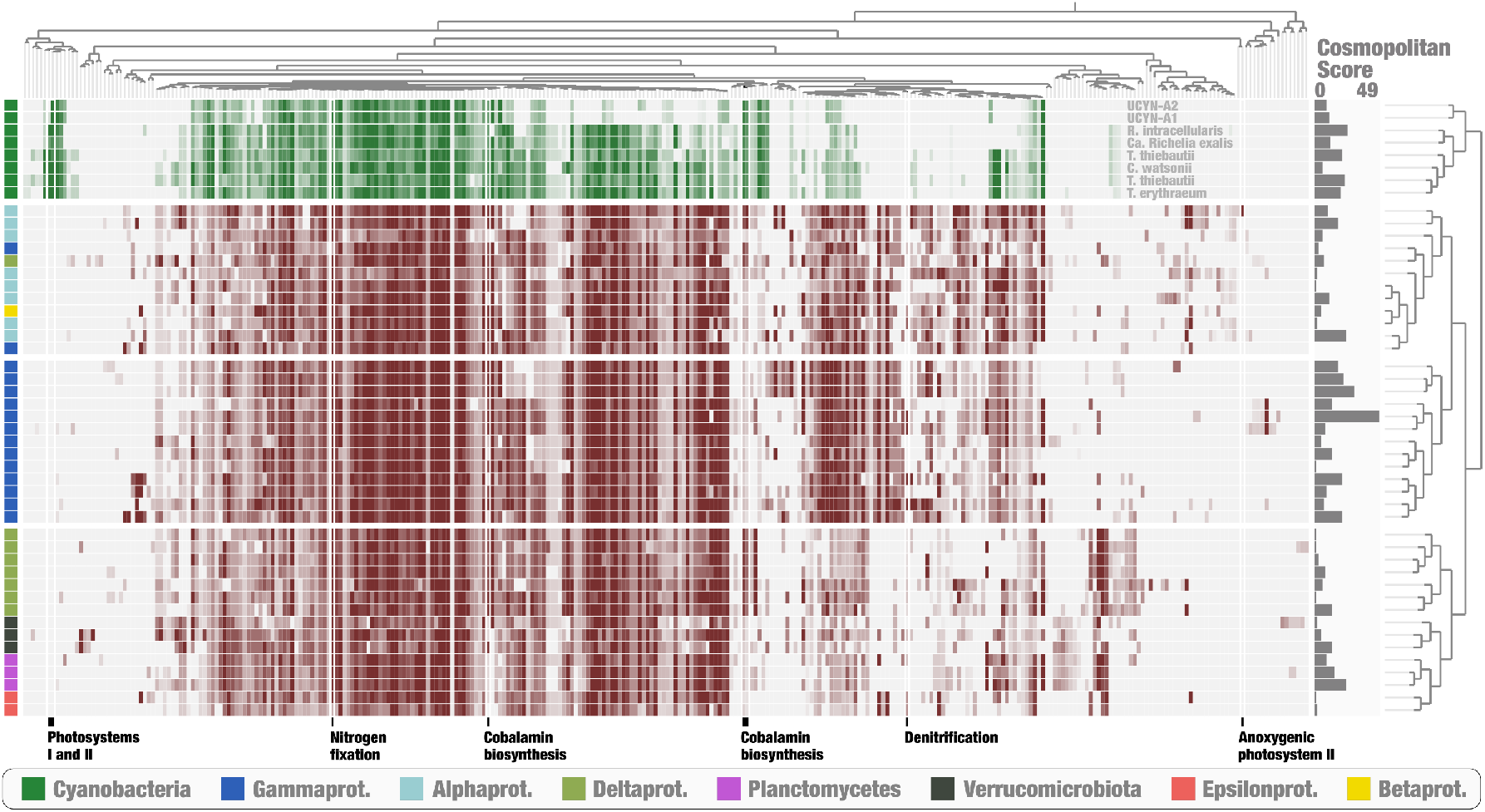
Functional life style of marine diazotrophs. The figure displays a heatmap of the completeness of 322 functional modules across the 48 diazotrophic MAGs. MAGs and modules were clustered based on the completeness values (Euclidean distance and ward linkage) and the data visualized using anvi’o^43^. The cosmopolitan score corresponds to the number of stations in which a given MAG was detected (cut-off: only when >25% of the MAG is covered by metagenomic reads).

#### HBDs are generally more abundant compared to cyanobacterial diazotrophs

The 48 diazotrophs were detected in up to 49 stations (out of 119 stations considered to compute this cosmopolitan score) and recruited up to 3.7% of metagenomic reads (Figures 1, 2 and Table S2) when considered individually. Yet, diazotrophs found to be most abundant locally were not the most widespread (R^2^ of 0.007 when comparing the maximal number of recruited reads and cosmopolitan score). In terms of geographical distributions, the diazotrophs remained undetected in the Arctic Ocean and a single HBD was detected in the Southern Ocean^39^. Furthermore, HBDs were undetected in the Red Sea and barely detected in the Mediterranean Sea. Within temperate and tropical regions of the open ocean, the Epsilonproteobacteria, Deltaproteobacteria and Verrucomicrobia marine diazotrophs were mostly detected in the Pacific Ocean. Remaining lineages occurred in the Pacific, Indian and Atlantic Oceans. It is worth mentioning that within cyanobacterial diazotrophs, the two populations of *Trichodesmium thiebautii*, highly abundant in some of the large size fractions, were mostly detected in the Indian Ocean (Figure 1). *Trichodesmium* might prevail in this region, but the overall geographic distribution of diazotrophs indicates that the Pacific Ocean is especially dominated by HBDs, corroborating previously observed trends^17,34,39^ with an extended set of diazotrophs and considering a wide size fraction for planktonic cells.

Among the 48 cyanobacterial and heterotrophic diazotrophs, 30 were mostly detected in the 0.2-5 μm size fraction covering most of the free-living bacterial cells, while the remaining diazotrophs were detected principally in the 5-20 μm (n=15) and 20-180 μm (n=2; *Richelia intracellularis* and ‘*Candidatus* Richelia exalis’) size fractions (Table S4). We then computed the ratio of cumulative mean coverage (i.e., number of times a genome is sequenced) between the eight cyanobacterial diazotrophs and 40 HBDs across 385 metagenomes organized by size fraction (552 metagenomes with no signal for any of the 48 diazotrophs were not considered here). Overall, HBDs displayed a cumulative mean coverage superior to that of cyanobacterial diazotrophs in 250 metagenomes, compared to 135 for the latter. Furthermore, a clear signal emerged in which HBDs were more abundant genome-wise compared to their cyanobacterial counterparts in most metagenomes from the 0.2-5 μm (86.5%) and 0.8-2000 μm (92.6%) size fractions while cyanobacterial diazotrophs predominated in the 20-180 μm (92.3%) and 180-2000 μm (86.2%) size fractions (Figure 1). Finally, the 5-20 μm size fraction was more balanced between HBDs and cyanobacterial diazotrophs.

The 0.8-2000 μm size fraction was unfortunately not collected during the first part of *Tara* Oceans sampling (specifically Mediterranean Sea, Red Sea and Indian Ocean), but provided a valuable metric to compare the relative abundance of diazotrophs in the Pacific and Atlantic Oceans that otherwise would be separated between the different size fractions. In other words, this size fraction could be used to effectively compare the genomic signal of diazotrophs corresponding to free-living, particle-attached, filamentous, colony-forming and symbiotic cells, provided they (or their hosts) pass filter holes 2 millimeter in diameter, either undamaged or fragmented (e.g., *Trichodesmium* colonies are known to be fragile). While uncertainty remains in the Indian Ocean, home to considerable *Trichodesmium* blooms based on data from *Tara* Oceans, trends from metagenomes corresponding to the 0.8-2000 μm size fraction in other regions largely mirrored the free-living size fraction since they were dominated typically by HBD signals. Critically, the 0.2-3 μm and 0.8-2000 μm size fractions indicate that HBDs are more abundant compared to their cyanobacterial counterparts in metagenomes covering most regions of the sunlit ocean.

#### Co-occurrence of HBDs in large size fractions from a Pacific Ocean station

We detected considerable metagenomic signal for HBDs at Station 98 in the South Pacific Ocean (Figure 3; Table S4), which was also found using reference *nifH* genes^17^. Station 98 includes five surface and three DCM metagenomes covering all size fractions except for 0.8-2000 μm. The only cyanobacterial diazotroph we detected in this metagenomic set was ‘*Candidatus* Richelia exalis’ with a mean coverage of just 0.4X in the 20-180 μm size fraction of the surface layer. The 40 HBDs remained undetected in the DCM and only two HBDs were slightly detected in the 0.2-3 μm size fraction of the surface layer. In marked contrast, 14 HBDs were detected in the 5-20 μm, 20-180 μm and 180-2000 μm size fractions of surface waters with a cumulative mean coverage reaching 1,106X (i.e., their genomes were sequenced cumulatively more than one thousand times in this particular metagenome), 15X and 283X, respectively. Such a high genomic coverage for bacterial populations in large size fractions is unusual and in the specific case of diazotrophs exceeded the highest metagenomic signal observed for UCYN-A and *Trichodesmium* in any oceanic region (Figure 3; Table S4). These 14 HBDs are affiliated to Deltaproteobacteria (n=5), Alphaproteobacteria (n=2), Gammaproteobacteria (n=2), Epsilonproteobacteria (n=2), Planctomycetes (n=2) and Verrucomicrobia (n=1). Samples collected at Station 98 contained very low concentrations of nitrate and this was especially true of the surface layer (0.001 μmol/L; Table S1). Thus, nitrogen depletion and other co-variables seemingly provided favorable conditions for a diverse assemblage of HBDs to occur abundantly in large size fractions of plankton. Lack of signal in the small size fraction suggests that similar populations might be missed in oceanic sampling that typically restrict bacterial analyses to free-living cells. Mechanisms maintaining diazotrophs in large plankton size fractions have yet to be fully elucidated^34,50–54^. Our results nonetheless echo recent observations in estuaries linking active HBDs to large aggregates that comprise polysaccharides^55^. Exopolymer particles and aggregate formations might create low-oxygen microenvironments for nitrogen fixation by HBDs in marine ecosystems^56^, as observed in culture conditions^57^. Thus, we suggest that HBDs formed a considerable number of large aggregates (up to >180 μm in size) at Station 98 in order to optimize their nitrogen fixation capabilities.

**Figure 3:**
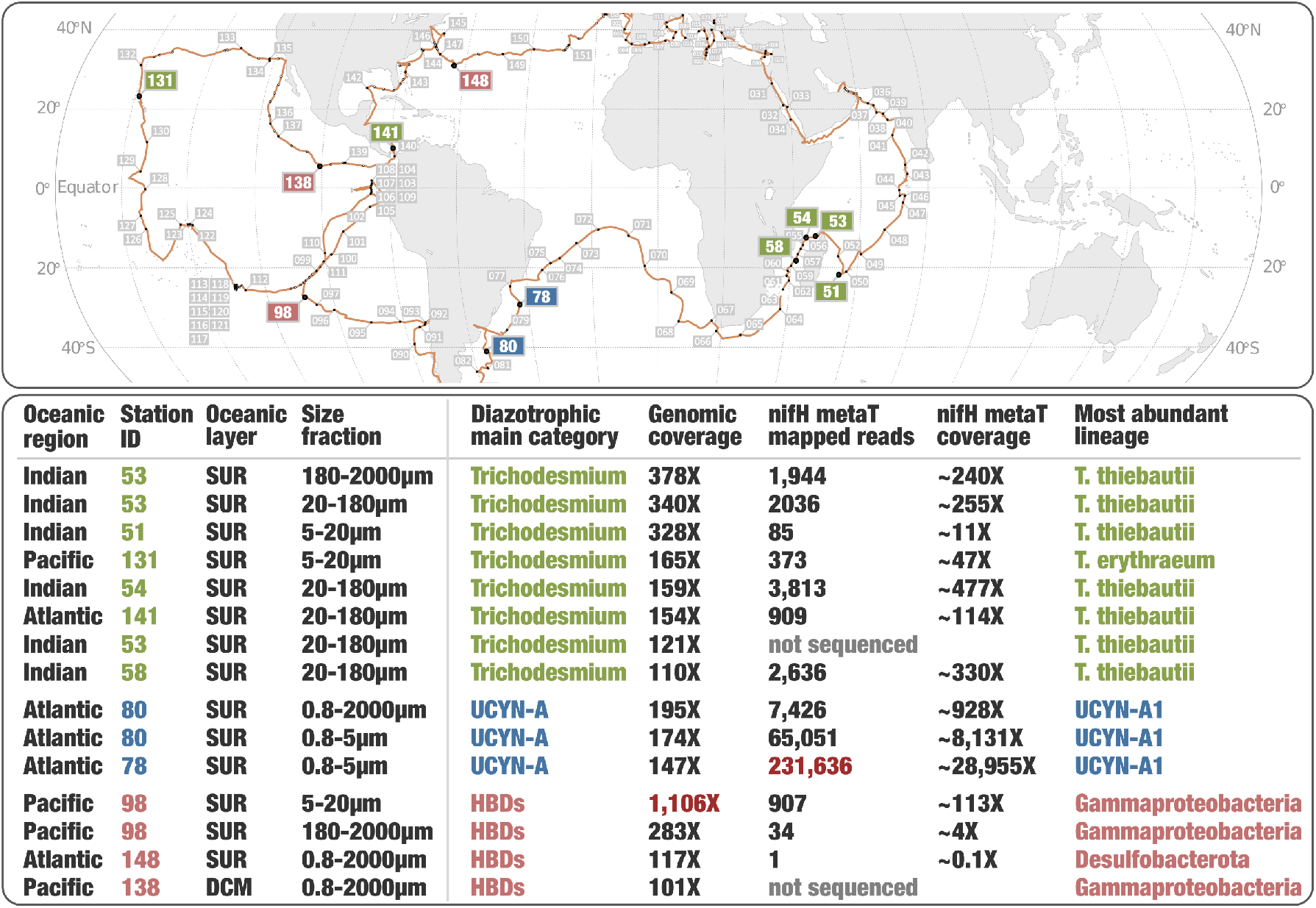
Oceanic stations with highest metagenomic signal for diazotrophs. The world map provides coordinates for 15 *Tara* Oceans metagenomes (10 stations) displaying cumulative genomic coverage >100X for MAGs affiliated to diazotrophic *Trichodesmium*, UCYN-A or the HBDs. Bottom panel summarizes multi-omic signal (including at the level of *nifH* genes) statistics for those 15 metagenomes.

### Part two: Gene-centric multi-omic analyses (*nifH* gene)

#### 48 diazotrophic MAGs may cover >90% of cells containing known *nifH* genes

In order to quantify the significance of 48 diazotrophic MAGs with regard to marine diazotrophs, we combined their *nifH* gene sequences with a comprehensive set of *nifH* sequences from culture, clones and amplicon surveys (see Methods) and used this extended *nifH* database (n=328; redundancy removal at 98% identity over 90% of the length) to recruit metagenomic reads from *Tara* Oceans (percent identity >90%; Table S8). Strikingly, *nifH* genes corresponding to the eight cyanobacterial diazotrophs and 40 HBDs recruited 42.3% and 49.1% of mapped metagenomic reads, respectively, with just 8.7% of the signal corresponding to 280 orphan *nifH* genes for which the genomic content within plankton has not yet been characterized (Figure 4 and Table S8). These include a well known diazotroph that awaits genomic characterization, the Gamma-A lineage^58^, which accounted for just 0.4% of mapped reads. Overall, this *nifH* centric metagenomic survey indicates that the 48 bacterial diazotrophic MAGs we have characterized account for more than 90% of cells containing known *nifH* genes and occurring in stations from the sunlit ocean sampled by *Tara* Oceans. One remaining uncertainty is the extent of abundant marine heterotrophic bacterial *nifH* genes that have yet to be discovered. These might further swell the ranks of HBDs in years to come.

**Figure 4:**
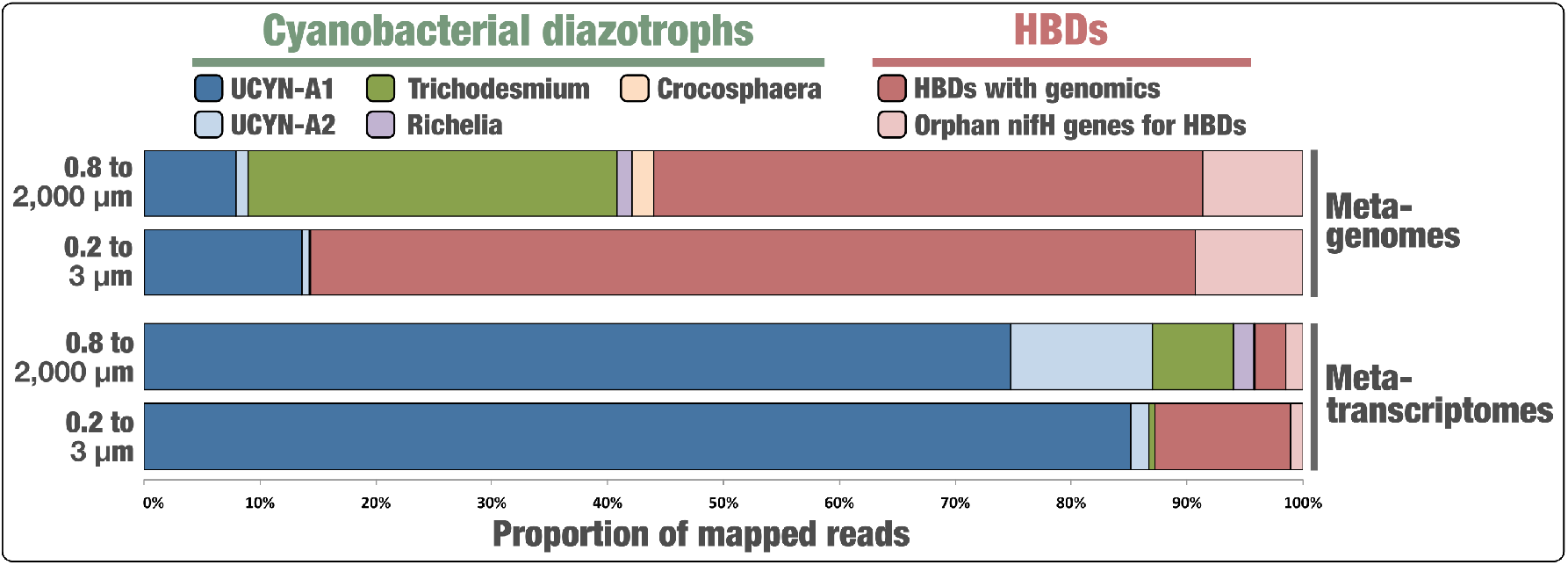
*nifH* gene detection across marine metagenomes and metatranscriptomes. The figure summarizes the proportion across diazotrophic lineages of mapped reads for the extended *nifH* database (328 sequences including 280 orphan genes) across *Tara* Oceans metagenomes (n=781) and metatranscriptomes (n=520) with signal and corresponding to surface and deep chlorophyll maximum layers. The 0.8-2000 μm layer summarizes signal for all size fractions within this range, including the 0.8-2000 μm size fraction.

#### HBD populations were found to express their *nifH* genes in the sunlit ocean

We mapped hundreds of *Tara* Oceans metatranscriptomes against the extended *nifH* database to gain some insights into the potential for nitrogen fixation activity of cyanobacterial diazotrophs and HBDs. Specifically, we recruited “bacteria-compatible” metatranscriptomic reads from the free-living size fraction (0.2-3 μm), as well as poly-A enriched metatranscriptomic reads from larger size fractions ranging from 0.8 μm to 2,000 μm that were produced primarily to explore the transcriptomic diversity of microbial eukaryotes^59^. Indeed, bacterial transcripts are not necessarily polyadenylated, and even when it does occur, polyadenylation is often a degradation signal^60^. Importantly, all of the HBD *nifH* genes recruited reads, indicating at the very least a basal expression of genes encoding the nitrogen fixation apparatus in every abundant marine diazotrophic lineage we have characterized with genomics thus far (Table S8). Furthermore, Station 98 with considerable genomic signal for HBDs also stood out with respect to metatranscriptomic signal for the corresponding *nifH* genes. Thus, HBDs are indeed expressing their *nifH* genes when found to be abundant under nitrate-depleted conditions of the Pacific Ocean.

When considering the extended *nifH* database as a whole, most of the signal among metatranscriptomes corresponded to UCYN-A1, followed by UCYN-A2, the HBDs and *Trichodesmium* (Figure 4 and Table S8). The predominance of UCYN-A signal (including in the “bacteria-enriched” 0.2-3 μm size fraction) was due to the apparently high nitrogen fixation activity for UCYN-A1 at Stations 78 and 80 in the South West region of the Atlantic Ocean in which hundreds of thousands of metatranscriptomic reads corresponded to its *nifH* gene alone (Figure 3), as already reported^61^. Metatranscriptomic read recruitments suggest that UCYN-A1 symbiont drives a substantial portion of the nitrogen fixation flux at the critical interface between oceans and the atmosphere, and this despite a relatively limited metagenomic signal (this genome was detected in just 13 stations). This metatranscriptomic analysis at large-scale substantiates the importance of UCYN-A as previously observed with *in situ* nitrogen fixation surveys (e.g.,^22^). However, the relatively low signal for *Trichodesmium* and HBDs was surprising. A trend emerged in which the *nifH* genes for symbiotic diazotrophs (UCYN-A, *Richelia*) were more significantly detected relative to their metagenomic signal compared to non-symbiotic diazotrophs, echoing other studies (e.g.,^62,63^). This could be due to symbiotic relationships favoring enhanced nitrogen fixation to the benefit of the diazotrophic hosts. However, one could also wonder if the hosts are protecting RNA molecules of bacterial origin during inherent sampling and filtration steps. Given that bacterial RNA molecules are highly unstable, marine metatranscriptomes should be interpreted with caution.

For now, the nitrogen fixation activity of HBDs versus cyanobacterial diazotrophs remains unclear. HBDs may contribute very little to nitrogen fixation rates among plankton, in particular as compared to UCYN-A, *Richelia*, and *Trichodesmium* populations. For instance, the streamlined genomes of UCYN-A populations and beneficial interactions with their hosts have created highly effective nitrogen fixation machineries^21,61,64^ compared to what HBDs can do by themselves and without ATP production from photosynthesis. Yet, metatranscriptomic surveys cannot be trusted to the same extent as metagenomes for semi-quantitative investigations, and do not equate to activity. Our only certitude at this point is that HBDs (1) are widespread and sufficiently abundant to make a real difference in the oceanic nitrogen balance, and (2) regularly transcribe their *nifH* gene in the sunlit ocean, including when co-occurring in large size fractions. These environmental genomic insights indicate that HBDs should not be excluded from the highly exclusive list of most relevant marine nitrogen fixers (currently only represented by cyanobacterial lineages^9^), at least until extensive studies of putative aggregates in the field as well as culture conditions shed light on their functional life style and metabolic activities.

#### A simple nomenclature to keep track of genome-resolved marine HBDs

As an effort to maintain some continuity between studies, here we suggest applying a simple nomenclature to name with a numerical system the non-redundant HBD MAGs with sufficient completion statistics as a function of their phylum-level affiliation (historic NCBI naming). For example, HBDs affiliated to Alphaproteobacteria and discovered thus far were named HBD Alpha 01 to HBD Alpha 08. Table S3 describes the 40 HBDs using this nomenclature, which could easily be expanded moving forward. To this point, only MAGs with completion >70% are part of this environmental genomic database, and the redundancy removal was set to ANI of 98%. Their genomic content can be accessed from https://figshare.com/articles/dataset/Marine_diazotrophs/14248283.

## Conclusion

Our genome-resolved metagenomic survey of plankton in the surface of five oceans and two seas covering organismal sizes ranging from 0.2 μm to 2,000 μm has allowed us to go beyond cultivation and *nifH* amplicon surveys to characterize the genomic content and geographic distribution of key diazotrophs in the ocean. Briefly, we identified eight cyanobacterial diazotrophs, seven of which were already known at the species level, and 40 HBDs, 32 of which were first characterized in this study. The 40 HBDs are functionally diverse and expand the known diversity of abundant marine nitrogen fixers within Proteobacteria and Planctomycetes while also covering Verrucomicrobia. Overall, the 48 diazotrophs characterized here may account for more than 90% of cells containing known *nifH* genes and occurring in the sunlit ocean. In other words, the genomic search for most abundant diazotrophs at the surface of the open ocean may be nearing completion.

Nitrogen fixers in the sunlit ocean have long been categorized into two main taxonomic groups: a few cyanobacterial diazotrophs contributing to most of the fixed nitrogen input^14,19,22,65^, and a wide range of non-cyanobacterial diazotrophs considered to have little impact on the marine nitrogen balance, in part due to their very low abundances within plankton as seen from several *nifH* based amplicon surveys^25–32^. Here we provide three results contrasting with this paradigm. First, we found that HBDs can occasionally co-occur under nitrate-depleted conditions in large size fractions, with metagenomic signals exceeding what was observed for UCYN-A and *Trichodesmium* lineages in other oceanic regions. Critically, insights from estuaries^55,57^ can explain the signal for HBDs in large size fractions of the open ocean, suggesting they form aggregates that provide low-oxygen microenvironments optimized for nitrogen fixation. These insights could explain, at least to some extent, high nitrogen fixation rates previously observed in parts of the Pacific Ocean that are depleted in cyanobacterial diazotrophs, which at the time was referred to as a paradox^45^. But most importantly, genome-wide metagenomic read recruitments for the 48 diazotrophs indicated that HBDs are more abundant than their cyanobacterial counterparts in most regions of the sunlit ocean. Metagenomes covering a wide size range for plankton (the 0.8-2000 μm size fraction) were critical to reach this conclusion. Mismatches between the widely used “nifH4” primer and *nifH* genes from most HBDs might explain to some extent the growing gap between prior *nifH* based sequence surveys and what genome-resolved metagenomics can reveal. Finally, we found that all HBDs express their *nifH* genes, including when co-occurring in large size fractions, expanding on previous observations based on a subset of the lineages in the 0.2-3 μm size fraction^40^. As a result, a new understanding is emerging from large-scale multi-omic surveys that depicts nitrogen fixers in the sunlit ocean as the sum of a few cyanobacterial diazotrophs together with a wider range of HBDs (more taxa and functions), all capable of using their nitrogen fixation gene machinery while thriving in specific size fractions and oceanic regions. Surveying HBD aggregates might represent a key new asset in understanding the marine nitrogen cycle.

Now that genome-resolved metagenomics has shed light on dozens of abundant marine HBDs, first within the limited scope of free-living cells^39^, and now by covering a much wider size range of plankton, it becomes apparent how little we know about their functional lifestyles in general, and role in oceanic primary productivity via nitrogen fixation rates in particular. Moving forward, it will be critical to enrich or cultivate these HBDs in the laboratory, as done for some of the key cyanobacterial diazotrophs decades ago^66^ or HBDs from estuaries more recently^57^. Culture conditions and dedicated *in situ* investigations will test whether or not HBDs can contribute more to nitrogen fixation rates compared to their cyanobacterial counterparts in large oceanic regions. This line of research should strongly benefit our understanding of nitrogen budgets in the open ocean.

## Material and methods

### *Tara* Oceans metagenomes

We analyzed a total of 937 *Tara* Oceans metagenomes available at the EBI under project PRJEB402 (https://www.ebi.ac.uk/ena/browser/view/PRJEB402). Table S1 reports general information (including the number of reads and environmental metadata) for each metagenome.

### Genome-resolved metagenomics

The 798 metagenomes corresponding to size fractions ranging from 0.8 μm to 2 mm were previously organized into 11 ‘metagenomic sets’ based upon their geographic coordinates^42^. Those 0.28 trillion reads were used as inputs for 11 metagenomic co-assemblies using MEGAHIT^67^ v1.1.1, and the scaffold header names were simplified in the resulting assembly outputs using anvi’o^43^ v.6.1. Co-assemblies yielded 78 million scaffolds longer than 1,000 nucleotides for a total volume of 150.7 Gbp. Here, we performed a combination of automatic and manual binning on each co-assembly output, focusing only on the 11.9 million scaffolds longer than 2,500 nucleotides, which resulted in 1,925 manually curated bacterial and archaeal metagenome-assembled genomes (MAGs) with a completion >70%. Briefly, (1) anvi’o profiled the scaffolds using Prodigal^68^ v2.6.3 with default parameters to identify an initial set of genes, and HMMER^69^ v3.1b2 to detect genes matching to bacterial and archaeal single-copy core gene markers, (2) we used a customized database including both NCBI’s NT database and METdb to infer the taxonomy of genes with a Last Common Ancestor strategy^59^ (results were imported as described in http://merenlab.org/2016/06/18/importing-taxonomy), (3) we mapped short reads from the metagenomic set to the scaffolds using BWA v0.7.15^70^ (minimum identity of 95%) and stored the recruited reads as BAM files using samtools^71^, (4) anvi’o profiled each BAM file to estimate the coverage and detection statistics of each scaffold, and combined mapping profiles into a merged profile database for each metagenomic set. We then clustered scaffolds with the automatic binning algorithm CONCOCT^72^ by constraining the number of clusters per metagenomic set to a number ranging from 50 to 400 depending on the set. Each CONCOCT clusters (n=2,550, ~12 million scaffolds) was manually binned using the anvi’o interactive interface. The interface considers the sequence composition, differential coverage, GC-content, and taxonomic signal of each scaffold. Finally, we individually refined each bacterial and archeal MAG with >70% completion as outlined in Delmont and Eren^73^, and renamed scaffolds they contained according to their MAG ID. Table S2 reports the genomic features (including completion and redundancy values) of the bacterial and archaeal MAGs.

### MAGs from the 0.2–3 μm size fraction

We incorporated into our database 673 bacterial and archaeal MAGs with completion >70% and characterized from the 0.2–3 μm size fraction^39^, providing a set of MAGs corresponding to bacterial and archaeal populations occurring in size fractions ranging from 0.2 μm to 2 mm.

### Characterization of a non-redundant database of SMAGs

We determined the average nucleotide identity (ANI) of each pair of MAGs using the dnadiff tool from the MUMmer package^74^ v.4.0b2. MAGs were considered redundant when their ANI was >98% (minimum alignment of >25% of the smaller SMAG in each comparison). We then selected the MAG with best statistics (highest value when computing completion minus redundancy) to represent a group of redundant MAGs. This analysis provided a non-redundant genomic database of 1,888 MAGs.

### Taxonomical inference of MAGs

We determined the taxonomy of MAGs using both ChekM^75^ and version 86^76^. However, we used NCBI taxonomy from the GTDB output to describe the phylum of MAGs in the results and discussion sections, in order to be in line with the literature.

### Biogeography of MAGs

We performed a final mapping of all metagenomes to calculate the mean coverage and detection of the MAGs. Briefly, we used BWA v0.7.15 (minimum identity of 90%) and a FASTA file containing the 1,888 non-redundant MAGs to recruit short reads from all 937 metagenomes. We considered MAGs were detected in a given filter when >25% of their length was covered by reads to minimize non-specific read recruitments^39^. The number of recruited reads below this cut-off was set to 0 before determining vertical coverage and percent of recruited reads.

### Cosmopolitan score

Using metagenomes from the Station subset 1 (n=757; excludes the 0.8-2000 μm size fraction lacking in the first leg of the *Tara* Oceans expeditions), MAGs were assigned a “cosmopolitan score” based on their detection across 119 stations, as previously quantified for eukaryotes^42^.

### Identification of diazotroph MAGs

In a first step, we used three HMM models from Pfam^77^ within anvi’o (e-value citoff of e-15) and targeting the catalytic genes (nifH, nifD, nifK) and biosynthetic genes (nifE, nifN, nifB) for nitrogen fixation. We then ran Interproscan^78^ on genes with a HMM hit and used TIGRFAMs^79^ results (we found those to the most relevant for nitrogen fixation) to identify diazotroph MAGs. Finally, we used RAST^80^ as a complementary approach to identify nitrogen fixing genes the HMM/Inteproscan approach failed to characterize. Among the 48 diazotroph MAGs, only one single gene (nifH) was not recovered with this approach. The most likely explanation is that the gene is simply missing from the MAG.

### Functional inferences of diazotroph MAGs

We inferred functions among the genes of diazotrophic MAGs using COG20 functions, categories and pathways^47^, KOfam^48^, KEGG modules and classes^49^ within the anvi’o genomic workflow^43^. Regarding the KOfam modules, we calculated their level of completeness in each genomic database using the anvi’o program “anvi-estimate-metabolism” with default parameters. The URL https://merenlab.org/m/anvi-estimate-metabolism describes this program in more detail.

### Sequence novelty for the *nifH* genes

The 47 *nifH* genes identified in the MAGs were considered novel if their sequence identity scores never exceeded 98% identity over an alignment of al least 200 nucleotides, when compared to a recently built *nifH* gene catalog by Pierella Karlusich et a.^17^ using blast^81^. Briefly, the *nifH* gene catalog consists of sequences from Zehr laboratory (mostly diazotroph isolates and environmental clone libraries; https://www.jzehrlab.com), sequenced genomes, and additional sequences retrieved from *Tara* Oceans metagenomic assemblies (co-assemblies^39^ and the OM-Reference Gene Catalog version 2^40^).

### A new database of *nifH* genes including diazotroph MAGs

We created a database of *nifH* genes covering the diazotroph MAGs as well as few hundred sequences from Pierella Karlusich et al.^17^ with signal in *Tara* Oceans metagenomes. We removed redundancy (cut-off=98% identity) between the diazotroph MAGs and the Pierella Karlusich database, except for *Trichodesmium thiebautii* due to the occurrence of multiple populations (and slight differences between MAGs and culture representatives) that stressed the need to further explore *nifH* gene microdiversity within this species. We performed a mapping of metagenomes and metatranscriptomes to calculate the mapped reads and mean coverage of sequences in the extended *nifH* gene database. Briefly, we used BWA v0.7.15 (minimum identity of 90%) and a FASTA file containing the sequences to recruit short reads.

### Phylogenetic analyses of diazotroph MAGs

We used PhyloSift^82^ v1.0.1 with default parameters to infer associations between MAGs in a phylogenomic context. Briefly, PhyloSift (1) identifies a set of 37 marker gene families in each genome, (2) concatenates the alignment of each marker gene family across genomes, and (3) computes a phylogenomic tree from the concatenated alignment using FastTree^83^ v2.1. We used anvi’o to visualize the phylogenomic tree in the context of additional information and root it at the level of to the phylum Cyanobacteria.

### Metatranscriptomic read recruitment for *nifH* genes

We performed a mapping of 587 *Tara* Oceans metatranscriptomes to calculate the mean coverage of sequences in the extended *nifH* gene database. Briefly, we used BWA v0.7.15 (minimum identity of 90%) and a FASTA file containing the nifH gene sequences to recruit short reads from all 587 metatranscriptomes.

## Data availability

All data our study generated are publicly available at http://www.genoscope.cns.fr/tara/ (metagenomic co-assemblies, FASTA files) or https://figshare.com/articles/dataset/Marine_diazotrophs/14248283 for the supplemental tables and information, as well as the genomic content of 48 marine diazotrophs using the new nomenclature (diazotrophic genomic database).

## Contributions

Tom O. Delmont conducted the study and performed the primary data analysis. Eric Pelletier and Juan Pierella Karlusich performed analyses regarding the abundance of MAGs and *nifH* genes (including helping creating the extended nifH gene database) across *Tara* Oceans metagenomes and metatranscriptomes. Iva Veseli and Jessika Fuessel performed functional analyses of the diazotrophic MAGs. A. Murat Eren computed to compatibility between *nifH* genes and widely used primers. All authors helped interpret the data. Tom O. Delmont wrote the manuscript, with critical inputs from all the authors.

## Acknowledgments

Our survey was made possible by two scientific endeavors: the sampling and sequencing efforts by the *Tara* Oceans Project, and the bioinformatics and visualization capabilities afforded by anvi’o. We are indebted to all who contributed to these efforts, as well as other open-source bioinformatics tools for their commitment to transparency and openness. *Tara* Oceans (which includes the *Tara* Oceans and *Tara* Oceans Polar Circle expeditions) would not exist without the leadership of the *Tara* Oceans Foundation and the continuous support of 23 institutes (https://oceans.taraexpeditions.org/). Some of the computations were performed using the platine, titane and curie HPC machine provided through GENCI grants (t2011076389, t2012076389, t2013036389, t2014036389, t2015036389 and t2016036389).

## Supplemental figures

**Figure S1:**
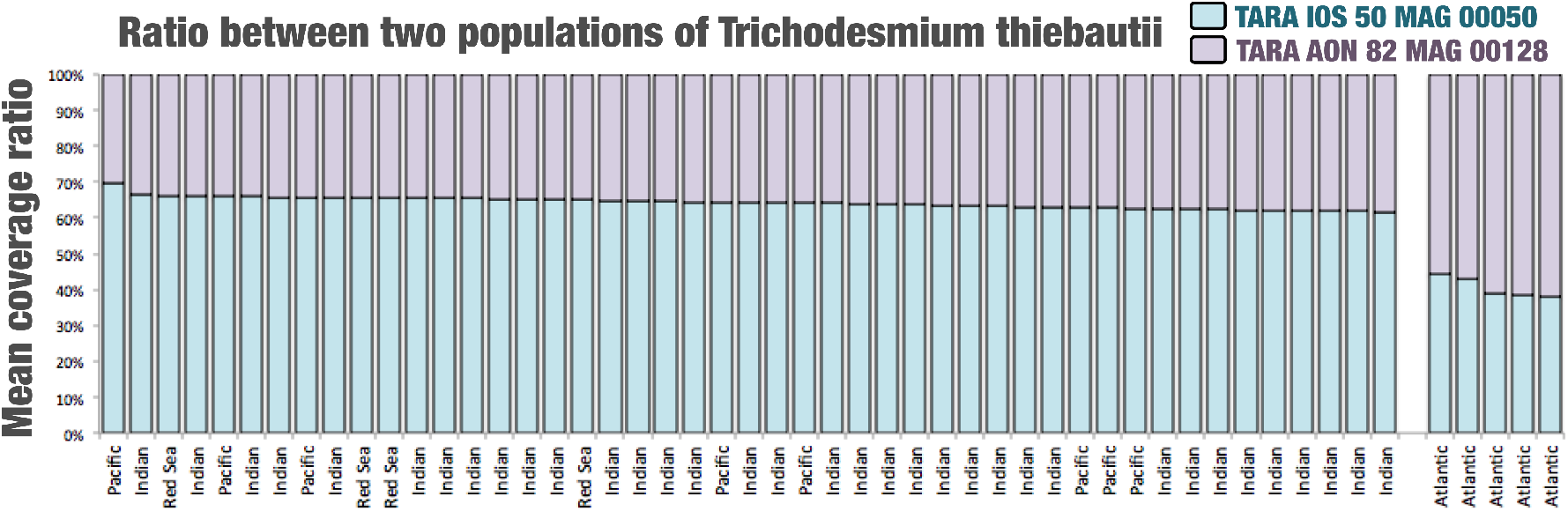
Mean coverage ratio for the two *Trichodesmium thiebautii* MAGs across 52 *Tara* Oceans metagenomes displaying a cumulative coverage >2X. Station Ids and associated data are available in the Table S4.

**Figure S2:**
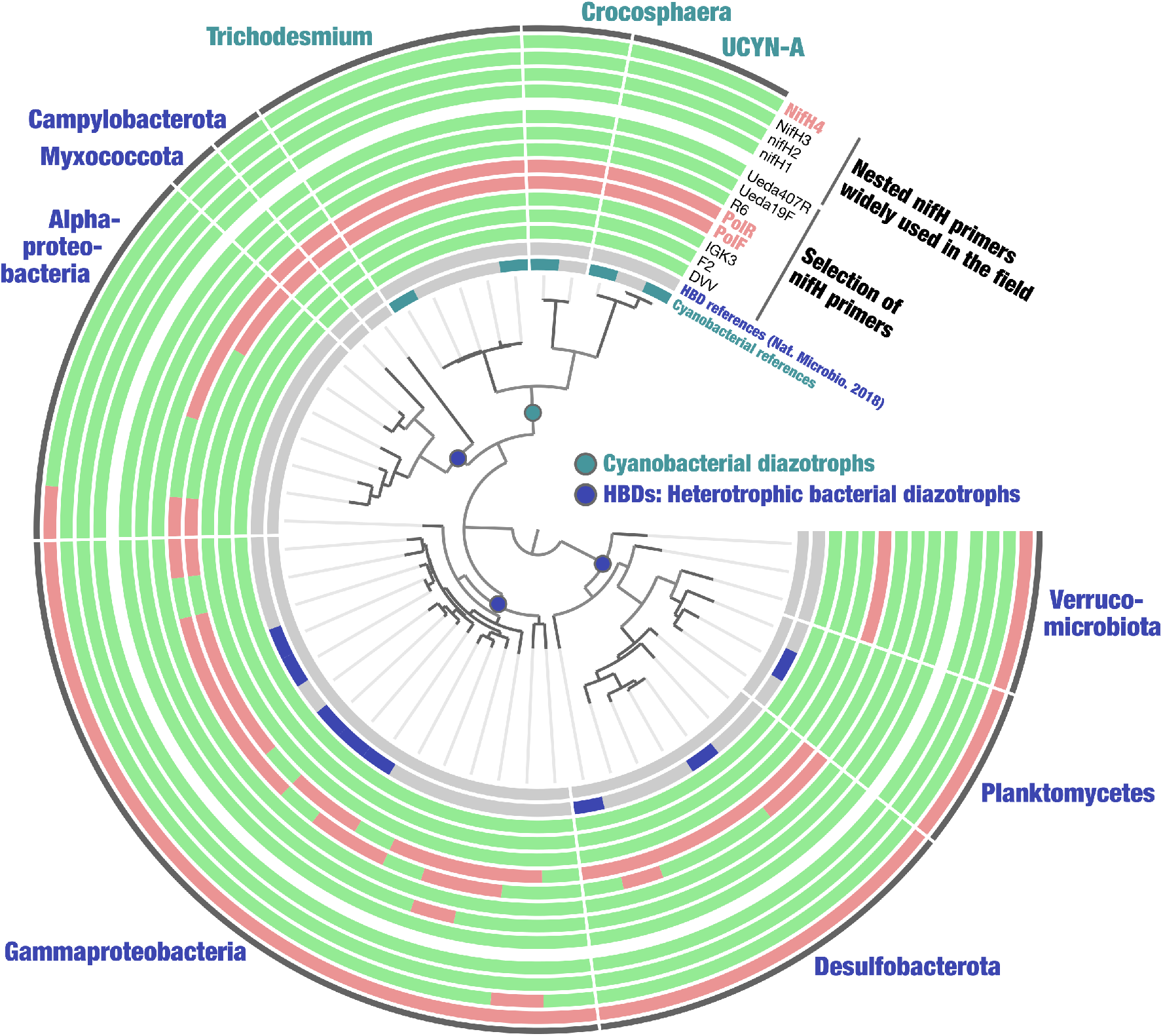
Interplay between the phylogeny and primer compatibility of *nifH* genes. The inner tree represents a phylogenetic tree of *nifH* genes from MAGs in our survey plus cyanobacterial references (built at the amino acid level with fastree^83^ within Genomenet (https://www.genome.jp/tools-bin/ete). Layers represent the compatibility (green) or incompatibility (red) of specific *nifH* primers used in the field (including for large-scale amplicon surveys).

## Supplemental Tables

**Table S01:** Statistics for 937 *Tara* Oceans metagenomes organized by depth, size fraction and oceanic region. The table also contains environmental conditions across *Tara* Oceans stations.

**Table S02:** Statistics for the 1,888 bacterial and archaeal MAGs. The table contains genomic statistics (e.g., completion and length), taxonomic information, general mapping trends such as the cosmopolitan score, as well as additional information regarding the 48 diazotroph MAGs.

**Table S03:** Occurrence of genes that encode the catalytic (nifHDK) and biosynthetic (nifENB) across 48 diazotrophic MAGs.

**Table S04:** Genome-wide metagenomic read recruitment statistics for the 48 diazotroph MAGs. The table contains mean genomic coverage values across the 937 *Tara* Oceans metagenomes described in Table S01.

**Table S05:** The *nifH* gene of 48 diazotroph MAGs. The table contains blast results when comparing *nifH* gene sequences from the diazotrophic MAGs to a reference *nifH* gene catalog.

**Table S06:** Compatibility between nifH primers and diazotrophic MAGs.

**Table S07:** Completeness of functional modules across the 48 diazotrophic MAGs.

**Table S08:** Metagenomic and metatranscriptomic mapping for the extended *nifH* database. The table also includes *nifH* gene sequences.

